# Radiosynthesis and preclinical evaluation of a carbon-11 labeled PDE7 inhibitor for PET neuroimaging

**DOI:** 10.1101/2021.06.12.447900

**Authors:** Zhiwei Xiao, Jiyun Sun, Masayuki Fujinaga, Huiyi Wei, Chunyu Zhao, Ahmed Haider, Richard Van, Tomoteru Yamasaki, Yiding Zhang, Jian Rong, Kuan Hu, Jiahui Chen, Erick Calderon Leon, Atsuto Hiraishi, Junjie Wei, Yi Xu, Yihan Shao, Han-Ting Zhang, Ying Xu, KC Kent Lloyd, Lu Wang, Ming-Rong Zhang, Steven Liang

## Abstract

**Background:** Dysfunction of cyclic nucleotide phosphodiesterase 7 (PDE7) has been associated with excess intracellular cAMP concentrations, fueling pathogenic processes that are implicated in neurodegenerative disorders. The aim of this study was to develop a suitable PDE7-targeted positron emission tomography (PET) probe that allows non-invasive mapping of PDE7 in the mammalian brain.

**Methods:** Based on a spiro cyclohexane-1,4’-quinazolinone scaffold with known inhibitory properties towards PDE7, we designed and synthesized a methoxy analog that was suitable for carbon-11 labeling. Radiosynthesis was conducted with the respective desmethyl precursor using [^11^C]MeI. The resulting PET probe, codenamed [^11^C]**26**, was evaluated by cell uptake studies, *ex vivo* biodistribution and radiometabolite studies, as well as *in vivo* PET experiments in rodents and nonhuman primates (NHP).

**Results:** Target compound **26** and the corresponding phenolic precursor were synthesized in 2-3 steps with overall yields of 49.5% and 12.4%, respectively. An inhibitory constant (IC_50_) of 31 nM towards PDE7 was obtained and no significant interaction with other PDE isoforms were observed. [^11^C]**26** was synthesized in high molar activities (170 - 220 GBq/µmol) with radiochemical yields of 34±7%. *In vitro* cell uptake of [^11^C]**26** was 6-7 folds higher in PDE7 overexpressing cells, as compared to the controls, whereas an *in vitro* specificity of up to 90% was measured. *Ex vivo* metabolite studies revealed a high fraction of intact parent in the rat brain (98% at 5 min and 75% at 30 min post injection). Considerable brain penetration was further corroborated by *ex vivo* biodistribution and PET imaging studies – the latter showing heterogenic brain uptake. While marginal specific binding was observed by PET studies in rodents, a moderate, but dose-dependent, blockade was observed in the NHP brain following pretreatment with non-radioactive **26**.

**Conclusion:** In this work, we report on the preclinical evaluation of [^11^C]**26** (codename [^11^C]P7-2104), a PDE7-targeted PET ligand that is based on a spiroquinazolinone scaffold. [^11^C]**26** displayed promising *in vitro* performance characteristics, a moderate degree of specific binding in PET studies with NHP. Accordingly, [^11^C]**26** will serve as a valuable lead compound for the development of a new arsenal of PDE7-targeted probes with potentially improved *in vivo* specificity.

## Introduction

The intracellular second messengers, cyclic adenosine monophosphate (cAMP) and cyclic guanosine monophosphate (cGMP), play a key role in signal transductions involving a myriad of processes in the central nervous system (CNS), immune system, as well as the cardiovascular system.^1-4^ Cyclic nucleotide phosphodiesterases (PDEs) constitute a superfamily of enzymes that degrade cAMP and cGMP. Human PDEs are derived from 21 genes and classified into 11 families (*PDE1-11*) according to the sequence homology of the *C*-terminal catalytic domain. Based on the substrate preferences, PDE families are grouped into cAMP-specific PDEs (4, 7 and 8), cGMP-specific PDEs (5, 6 and 9), and dual cAMP/cGMP-specific PDEs (1-3, 10 and 11).^5, 6^ Among the cAMP-specific PDEs, PDE7 has recently gained increasing attention due to the broad implications in pro-inflammatory processes that involve T cell activation. ^7^ Of note, the PDE7 family can be sub-divided into PDE7A and PDE7B subunits.^8,9^ In terms of organ distribution, PDE7 was found to be primarily expressed in the brain and heart, followed by liver, skeletal muscles, kidneys, testes, and pancreas. In particular, PDE7B is highly expressed in the brain in which the highest mRNA level found in the striatum and low in the cerebellum.^9^ On a cellular level, PDE7 isozymes are abundant in a variety of inflammatory and immune cells.^9-12^ It is worth mentioning that the increased PDE7B expression was found in chronic lymphocytic leukemia cells.^13^

Intracellular cAMP levels are highly regulated and typically maintained in a narrow range for normal physiological functions. Indeed, PDE7 is considered crucial in maintaining homeostatic cAMP levels. Similarly, inhibition of PDE7 emerged as a promising therapeutic strategy for a variety of neurological, inflammatory and immunological disorders where intracellular cAMP concentration is perturbed.^14, 15^ A growing body of evidence suggested that inhibition of PDE7 might provide neuroprotective effects and be beneficial to the treatment of neurological disorders, especially neurodegenerative diseases, such as Alzheimer’s disease (AD),^16, 17^ Parkinson’s disease (PD)^18^ and multiple sclerosis (MS)^19^. Because the second messenger, cAMP, not only mediates the signal transduction of the intracellular inflammatory mediators, but also involves lymphocyte proliferation and the immune response with respect to cytokine/chemokine production,^20, 21^ PDE7 inhibitors harbor the potential for the treatment of inflammatory conditions, such as autoimmune diseases (AIDs), autoimmune hepatitis (AIH) and rheumatoid arthritis (RA).^15, 22-26^

In view of the potential of PDE7 as a drug target in neurological, inflammatory and immunological disorders, the development of PDE7 inhibitors has progressed well in recent years (Figure 1). In terms of chemical structure, heterocyclic compounds, which are summarized in previous reviews,^27, 28^ have received considerable attention. Indeed, quinazolinone, quinazolinethione^29^ and purine-2,6-dione derivatives^20, 30^ with PDE7 inhibitory properties have been developed and evaluated. In addition, dual-selective PDE4/PDE7,^20, 31-33^ PDE7/PDE8 and PDE7/GSK3 inhibitors^28^ were synthesized and assessed in animal models of neurological and peripheral inflammatory diseases. To the best of our knowledge, no PDE7-selective inhibitors are currently on the market or in clinical trials.^1^ Given that PDE7 shares a high degree of sequence homology with PDE4, most PDE7 inhibitors exert pharmacological activity against PDE4. Thus, the development of selective PDE7 inhibitors remains challenging.

**Figure 1.**
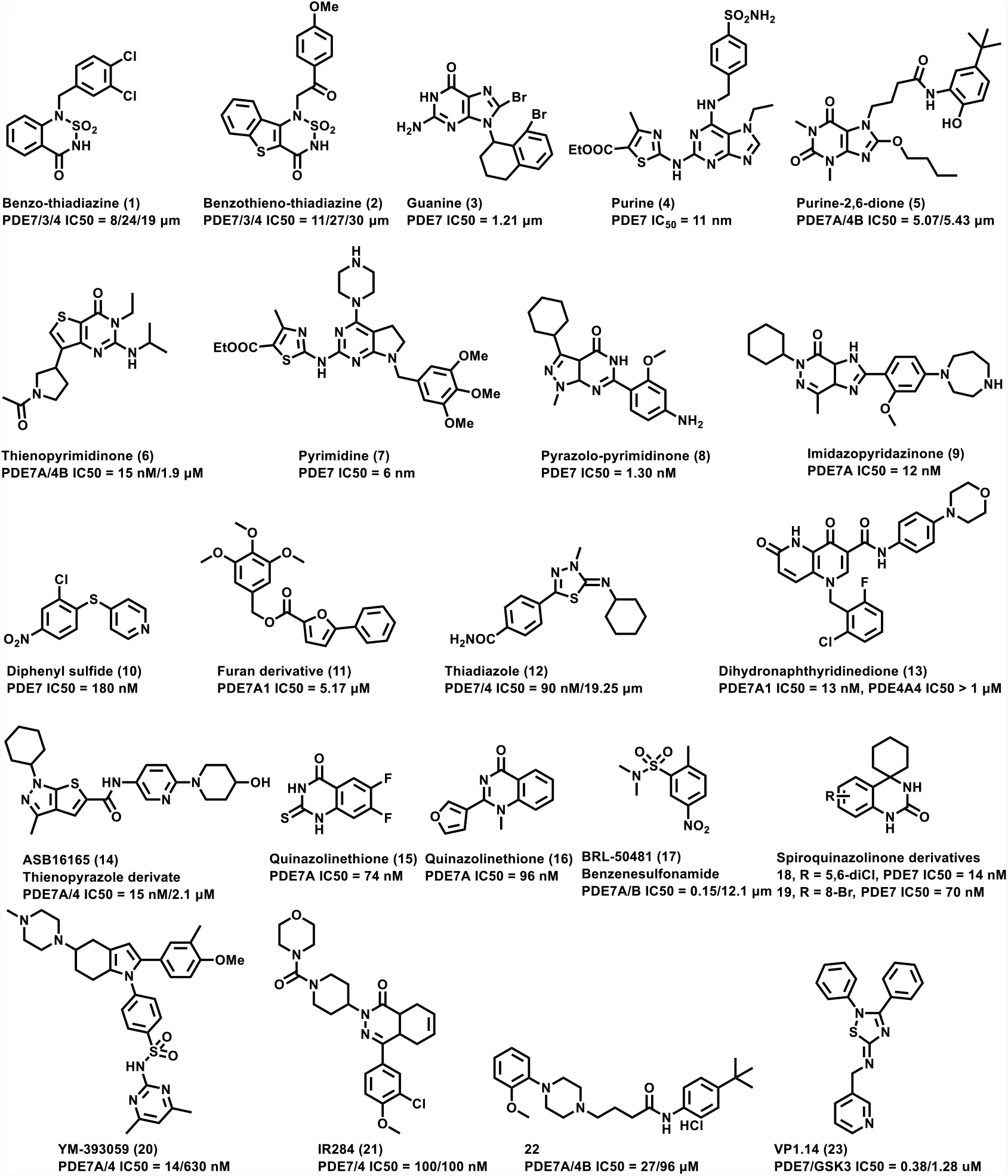
Selected compounds with PDE7 inhibitory properties.

Positron Emission Tomography (PET) is a non-invasive imaging modality widely used in preclinical and clinical settings due to its high sensitivity. As such, PET is ideally suited for early disease diagnosis, thereby providing prognostic information and allowing therapy monitoring. Further, PET can provide pharmacokinetic and pharmacodynamic information, which is crucial in early drug development to avoid late-stage failure of drug candidates.^34, 35^ The potential of PDE7 as a therapeutic target has been elucidated for several years,^36^ however, only a few PDE7 radioligands for PET neuroimaging have been reported to date. Recently, a PDE7-targeted PET radioligand, codenamed [^11^C]MTP38 (Figure 2), was advanced to humans.^37^ In the preclinical PET imaging study of rats and monkeys, [^11^C]MTP38 penetrated the BBB and rapidly accumulated in the striatum, while a lower tracer uptake was found in the cerebellum.^38^ Although only moderate in vivo binding specificity was observed, potentially attributed to relatively fast clearance, [^11^C]MTP38 served as a lead PET ligand scaffold to image PDE7 in the brain. Thomae *et al*. performed the synthesis and evaluation of [^18^F]MICA-003 in mice (Figure 2).^39^ Although MICA-003 exhibited a high affinity to PDE7 (17 nM) and readily crosses the blood-brain barrier, the rapid degradation to 2-[^18^F]fluoroethanol^40^ hampered its further development. Our study hypothesized that the substitution of metabolically labile fluoroethyl group in the chemical structure of MICA-003 by a methyl group would eliminate the generation of [^18^F]fluoroethanol in vivo and provide a promising concept for the development of a suitable PDE7-targeted PET ligand. Herein, we described the synthesis and preliminary pharmacology and ADME evaluation of compound **26** (codename P7-2104; Figure 2), followed by ^11^C-isotopologue labeling and PET imaging in rodents and nonhuman primates (NHPs), the results of which might provide an excellent chemical phenotype in medicinal chemistry aimed for further PDE7 PET ligand development.

**Figure 2.**
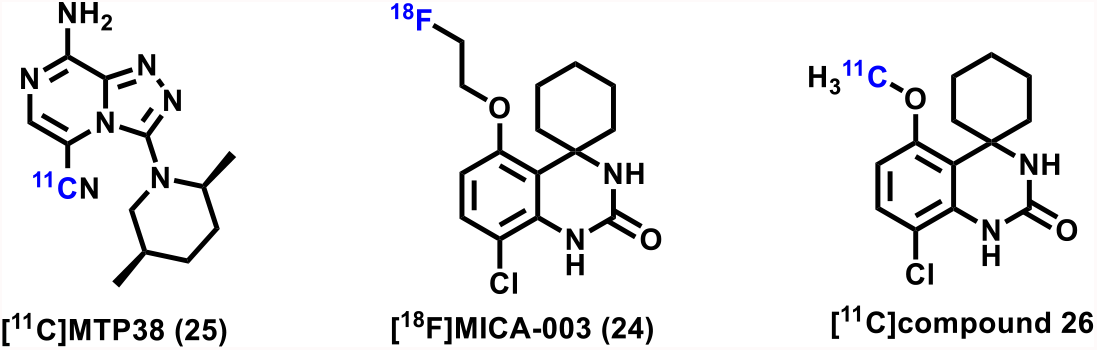
Reported PDE7 PET radioligands.

## Materials and methods

### Chemistry

Target compound **26** and the corresponding demethylated precursor **27** were synthesized as a previous publication,^41^ however, with minor modification, as shown in **Scheme 1**. Initial condensation between 2-chloro-5-methoxyaniline (**28**) and potassium isocyanate was accomplished under acidic conditions. In the presence of phosphorus pentoxide and methanesulfonic acid, intermediate **29** was reacted with cyclohexanone **30** to afford target compound **26** in 50% yield (over two steps). Precursor **27** was obtained through demethylation of compound **26** using hydrobromic acid at a high temperature.

**Scheme 1.**
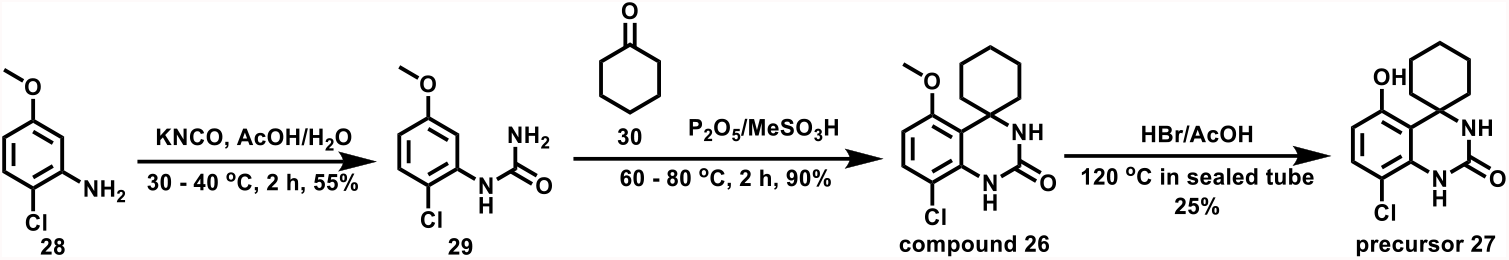
Synthesis of compound **26** and the corresponding precursor **27** for ^11^C labeling.

### Molecular Docking

The structure of compound **26** was generated using ChemDraw and IQmol. Since no crystal structure of PDE7B was available, a homology model was constructed using Swiss-Model.^42, 43^ The sequence of PDE7B was obtained from universal protein resource (uniprot.org) and a template structure (PDB 3G3N) was identified. After obtaining both the protein and ligand structures, they were docked globally by having the search box encompass the entire PDE7B protein and using AutoDock Vina in UCSF Chimera. The results for the protein-ligand complex were saved and were then loaded into Schrödinger Maestro to generate the ligand interaction plot.

### *In vitro* inhibitory assay to PDE7 of compound 26 and selectivity to other PDEs

Functional activity of target compound **26** for inhibition of PDE7A activity and the selectivity over other PDE isozymes (PDE1A/B/C, 2A, 3A/B, 4A/B/C/D, 5A, 8A, 9A, and 10A) were evaluated by Eurofins Panlabs Discovery Services and Reaction Biology Corp., respectively. Human recombinant enzymes were used for the assays. The inhibitory assays were performed and calculated in duplicate at 3 µM. If the inhibition percentage was less than 50%, the assay for half-maximal inhibitory concentration (IC_50_) value was conducted in duplicate.

### CYP450 isoform metabolism and *in vitro* safety profiling

The inhibitory constant of compound **26** toward cytochrome P450-dependent metabolic pathways was determined for individual CYP isoform (CYP1A2, CYP2C9, CYP2C19, CYP2D6, and CYP3A4). Reference inhibitors were included as positive controls. A 10 µM solution of compound **26** was used for initial screening. If the inhibition percentage was less than 50%, the experiment was stopped. If the inhibition potency was greater than 50%, the gradient concentrations of compound **26** were used for IC_50_ determination. The P450 reactions were detected at Ext/Em: 405 nm/460 nm (530 nm/605 nm for CYP2C9) in a time-dependent manner for 80 min with an acquisition interval of 2 min. IC_50_ curves were plotted using a non-linear regression model with Graphpad Prism 6. The latter method was further applied to assess the inhibitory constant of compound **26** to the hERG channel.

### Pharmacokinetics in Plasma and Brain (NeuroPK study)

Nine male wildtype Sprague Dawley rats were administered a bolus injection containing compound **26** in saline (1 mg/kg, 2 mL/kg) via the tail vein. Blood samples (approximately 120 µL) were harvested under light isoflurane anesthesia from a set of three rats at 0.08, 0.25, and 1 hr. Blood and brain samples were collected at 5, 25 and 60 min. Following centrifugation of blood samples, the resulting plasma supernatant was stored at −70 ± 10 °C till further analysis. Brain samples were homogenized using ice-cold phosphate buffer saline (pH 7.4) and homogenates were stored below −70 ± 10 °C until further analysis. Total homogenate volume was three times the brain weight. Plasma and brain samples were quantified by LC-MS/MS method. The plasma and brain concentration-time data of compound **26** was used for pharmacokinetic analysis.

### Radiochemistry

Carbon-11 labeling of compound **26** was achieved by radiomethylation of phenolic precursor **27** in the presence of potassium carbonate. The synthetic process involving radiolabeling, purification and formulation, was conducted using an automated module in an overall synthesis time of 60 min from the end of bombardment. [^11^C]CH_3_I was synthesized from cyclotron-produced [^11^C]CO_2_, which was obtained via the ^14^N(*p,α*)^11^C nuclear reaction. Briefly, [^11^C]CO_2_ was bubbled into a solution of LiAlH_4_ (0.4 M in THF, 300 μL). After evaporation, the resulting reaction mixture was treated with hydroiodic acid (57% aqueous solution, 300 μL). [^11^C]CH_3_I was transferred under helium gas with heating into a reaction vessel containing a solution of the precursor (1.0 mg) in anhydrous dimethyl sulfoxide (DMSO, 300 μL) with 1.0 mg K_2_CO_3_. After the radioactivity reached a plateau during the transfer, the reaction vessel was warmed to 100 °C and maintained for 5 min. The mobile phase (2.5 mL) and H_2_O (1.5 mL) were added to the reaction mixture, which was then injected into a semi-preparative HPLC system. HPLC purification was completed on a Phenomenex Luna 5μm C18 column,10 mm i.d. × 250 mm, UV at 254 nm, CH_3_CN/H_2_O = 50/50, 0.1%Et_3_N, flow = 5 mL/min. The radioactive fraction corresponding to the desired product was collected in a sterile flask, diluted with 30 mL of water, and trapped on a Sep-Pak light C18 cartridge. After washing with 10 mL of water, the product was eluted from the C18 cartridge with 0.3 mL of ethanol and formulated with 6 mL of saline. The radiochemical purities and molar activity were measured by analytical HPLC (Xselect Hss T3, 4.6 mm i.d. × 150 mm, UV at 254 nm, CH_3_CN/H_2_O = 50/50, 0.1% Et_3_N, flow = 1 mL/min). The identity of [^11^C]**26** was confirmed by co-injection with the unlabeled standard.

### Lipophilicity

The general procedure for lipophilicity measurement was previously described, ^44^ with minor modification in this work. Briefly, the measurement of Log*D* value was carried out by mixing [^11^C]**26** (radiochemical purity > 99%) with *n*-octanol (3.0 g) and PBS (0.1 M, 3.0 g) in a test tube. Both *n*-octanol and PBS were pre-saturated with each other prior to use. The tube was first vortexed for 5 min, followed by centrifugation (∼3500-4000 rpm) for an additional 5 min. PBS and *n*-octanol were aliquoted, weighed and the radioactivity in each component was measured using a Cobra Model 5002/5003 gamma counter. The Log*D* was determined by Log [ratio of radioactivity in *n*-octanol and aqueous layer, respectively] (n = 3).

### Plasma protein binding

For the evaluation of plasma protein binding,^45^ 55.5 MBq radiotracer was added to 150 μL of plasma, which was pre-incubated under 37 ^°^C for 5 min (n = 3). The samples were incubated at 37 ^°^C for 10 min. To each 150 μL of radiotracer-plasma solution was added 300 μL of ice-cold PBS and all samples were briefly vortexed. The samples were centrifuged at 14,000 g in Amicon centrifugal filters with a size cutoff of 10 kDa for 15 min at 4 °C, and the protein fraction was subsequently washed with 300 μL cold PBS at 21,000 g for 20 min at 4 °C. Additional cold PBS (300 µL) was used to wash the tube and collect all the filtrates. The radioactivity (in Becquerel) was measured in the protein fraction (A_protein_), filtrate, and filter unit using a gamma counter (Wizard, PerkinElmer), A_total_ was calculated as the sum of radioactivity, and the free fraction *f*_*u*_ was calculated according to the following eq.: *f*^*u*^ = 1-*A*^*protein*^*/A*^*total*^

### *In vitro* cell uptake

Control study: HEK293-PDE7B Recombinant cells (catalog#60412, BPS Bioscience) and HEK293 control cells (catalog# CRL-1573, ATCC) – both in logarithmic growth phase – were plated in a 24 well plate (2 × 10^5^ cells per well) and cultured overnight. After 24 hr, each well was added 2 μCi of [^11^C]**26**, and incubated at 37 ^°^C for 30 min or 60 min. The radioactive medium was aspirated and collected in a tube. The residual cells were washed twice with PBS, and then collected in the same tube. The cells were lysed with 1 N NaOH (200 μL) and washed twice with PBS, all the resulting solutions were collected in one tube. The radioactivity in the collected supernatant and cell lysis buffer (600 μL/tube) was measured using a Cobra Model 5002/5003 gamma counter, respectively.

Blocking study: HEK293-PDE7B cells in logarithmic growth phase were plated in 24 well plate (2 × 10^5^ cells per well) and cultured overnight. After 24 h, each well was added 2 μCi of [^11^C]**26** with or without unlabeled reference compound **26** or BRL50481, respectively. The resulting culture medium contained 2 μCi of [^11^C]**26**, 1 µM of unlabeled compound **26** or BRL50481, and 5% DMSO. Cells were incubated at 37 ^°^C for 60 min and samples were collected as described for the control study. The cell uptake was calculated according to the following eq.: *cell uptake%* = *A*_*lysis buffer*_*/*(*A*_*lysis buffer+*_*A*_*supernate*_) (n = 4)

### PET imaging in rodents

All animal experiments were approved by the Institutional Animal Care and Use Committee of Massachusetts General Hospital or the National Institute of Radiological Sciences (Japan). C57BL/6 mice (female; 8 weeks, 20–25 g) and Sprague-Dawley rats (male; 8-9 weeks; 264-310 g) were kept on a 12 h light/12 h dark cycle and were allowed food and water *ad libitum*.

Rodent PET scans were carried out using a Genisys 4 PET (Sofie Biosciences, Culver, CA, USA) or a Siemens Inveon PET/CT system. Animals were kept under anesthesia using 1-2% (v/v) isoflurane in oxygen during the scan. The radiotracer (1.85 MBq for mice; 42-53 MBq for rats) was injected into the tail vein via a preinstalled catheter. A dynamic scan in 3D list mode was acquired for 60 min. The PET dynamic images were reconstructed using the manufacturer’s acquisition software. Volumes of interest, including whole brain, cortex, striatum, hippocampus, thalamus and cerebellum were placed using AMIDE / PMOD software. The radioactivity was decay-corrected and expressed as standardized uptake value (SUV = radioactivity per mL tissue / injected radioactivity × bodyweight).

### PET imaging in nonhuman primates

Cynomolgus monkeys used for PET/CT scan were deprived of food for 12h but allowed to drink water at any time. Animals (Male, bodyweight 5.0-5.2 kg) were anesthetized with ketamine, placed into the scanner (GE Discovery Elite 690, USA), and maintained with 2% isoflurane and 98% oxygen. A solution of [^11^C]**26** (6.24–9.84 mCi) was injected into the monkey via a venous catheter, followed by a dynamic PET scan. For the blocking study, the blocking reagent reference compound **26** (0.4 or 1.0 mg/kg, iv) was used, followed by the injection of [^11^C]**26**. Time-activity curves of each brain region were extracted from the corresponding VOIs and brain uptake was decay-corrected and expressed as SUV.

### Whole-body *ex vivo* biodistribution studies in mice

A solution of [^11^C]**26** (1.85 MBq/100 µL) was injected into CD-1 mice via the tail vein. Animals were sacrificed at 5, 15, 30, and 60 min post radiotracer injection (n= 4 for each time point). Major organs, including the whole brain, heart, liver, lungs, spleen, kidneys, small intestine (including contents), muscle, pancreas, stomach, bone, and blood were quickly harvested and weighed. The radioactivity in these tissues was measured using a Cobra Model 5002/5003 gamma counter, and all radioactivity measurements were decay corrected based on the half-life of carbon-11. The results are expressed as the percentage of injected dose per gram of wet tissue (%ID/g).

### Radiometabolite Analysis

After intravenous injection of [^11^C]**26** through the tail vein, SD rats were sacrificed at 5 min and 30 min post injection. Blood samples were collected and centrifuged at 14,000 rpm for 3 min at 4 °C to separate the plasma. The supernatant was collected and added to an ice-cooled test tube containing 100 μL of CH_3_CN. After vortex for 10 s, the mixture was centrifuged at 14000 rpm for 3 min at 4 °C for deproteinization. The supernatant was collected, and the process was repeated until no precipitations were observed when CH_3_CN was added. The rat brain was immediately dissected, homogenized with 400 μL of ice-cooled CH_3_CN, and then centrifuged at 14,000 rpm for 5 min at 4 °C. The supernatant was collected in a test tube containing 100 μL of ice-cooled CH_3_CN, and the process (vortex and centrifuge) was repeated until no precipitations were observed when CH_3_CN was added. An aliquot of the supernatant (100 µL), obtained from the brain homogenate, was injected into and analyzed by a radio-HPLC system. The percentage of [^11^C]**26** to total radioactivity (corrected for decay) on the HPLC charts was calculated as (peak area for [^11^C]**26**/total peak area) × 100. The same procedure was used for metabolite analysis in plasma.

### Statistical analysis

Statistical analysis was performed by Student’s two-tailed *t-*test, and asterisks were used to indicate statistical significance: ^*^ *p* < 0.05, ^**^ *p* ≤ 0.01, ^***^ *p* ≤ 0.001 and ^****^*p* < 0.0001.

## Results

### Chemistry

Target compound **26** and the corresponding phenolic precursor **27** were efficiently synthesized in 2-3 steps from commercially available 2-chloro-5-methoxyaniline (**28**) with 49.5% and 12.4% overall yields (Scheme 1), respectively.

### *In vitro* inhibitory assay to PDE7 of compound 26 and selectivity to other PDEs

Compound **26** discovered by Pfizer Research,^41^ exhibited equipotency to PDE7 subtypes, namely PDE7A and PDE7B. Therefore, we only measured PDE7A to confirm the binding potency. In specific, by in vitro assessment, compound **26** exhibited an inhibitory constant (IC_50_) of 31 nM to PDE7A (Figure 3). Further, the target compound was found to be selective over other PDE isomers (% inhibition < 50% at 3 μM of compound **26**).

**Figure 3.**
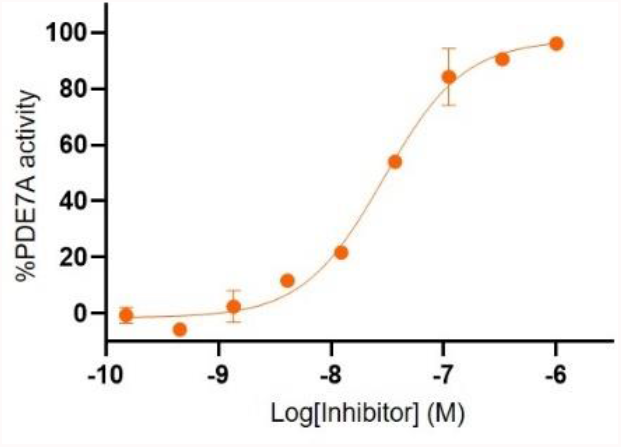
Concentration-response curve of compound **26** for inhibition of PDE7A activity.

### Molecular docking

Molecular docking was performed with a homology model of PDE7B to explore potential interactions. The binding pocket is mainly hydrophobic and major ligand-protein interactions were observed with Tyr 172, Phe345, and Phe377 (Figure 4A). Indeed, target compound **26** formed a hydrogen bond of 2.306 Å with Tyr172. Further, Pi-Pi stacking was observed between the ligand and Phe345/377 (Figure 4B).

**Figure 4.**
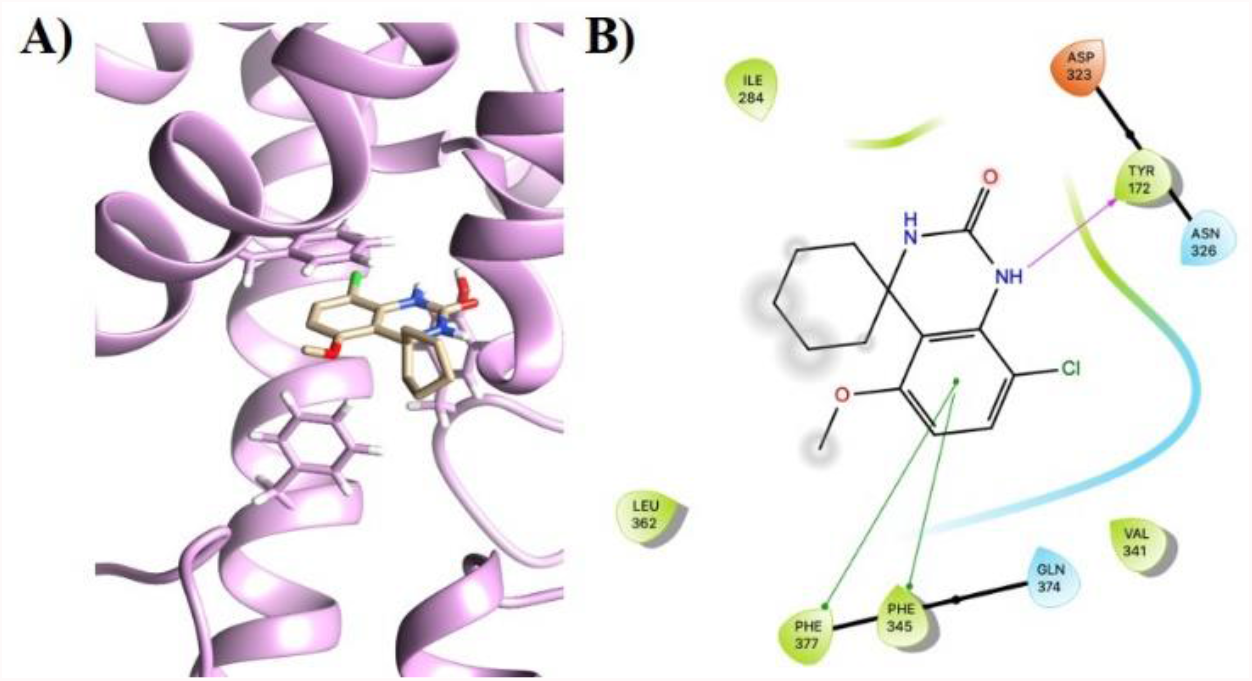
The docking position of compound **26** was generated using AutoDock Vina module of Chimera and Maestro. (A) AutoDock Vina (Chimera) generated model of compound **26** and (B) The corresponding Maestro generated interaction plot, where purple arrow represents a hydrogen bond and green arrows highlight Pi-Pi stacking.

### CYP450 isoform metabolism and in vitro safety profiling

By *in vitro* safety studies, compound **26** did not exert an inhibitory effect on the hERG channel (IC_50_ > 100 μM). Except for the weak inhibitory activity against CYP1A2 (IC_50_ = 4.4 μM), compound **26** did not markedly interact with any CYP450 isoforms (IC_50_ ≥ 10 μM).

### Pharmacokinetics in Plasma and Brain (NeuroPK study)

The brain-to-plasma ratio (Kp) was in the range of 3-5 at all measured time points, indicating that the concentration of compound **26** was higher in the brain than in the plasma throughout the study (Table 1 and Figure 5). While initially peaking at 3316.17 ± 378.14 ng/g (5 min), a steady washout was observed at later time points, suggesting fast and reversible kinetics. Indeed, less than 5% of compound remained in the brain at the last measured time point (60 min). A similar trend was found in the plasma, from 712.49 ± 153.23 ng/mL (5 min) to 39.39 ± 4.44 ng/mL (60 min).

**Table 1.** Rats PK data in plasma and brain for compound **26** (n =3)

| Time/min | Plasma Concentration (ng/mL) |  | Brain concentration (ng/g) |  | Brain -Kp |  |
| --- | --- | --- | --- | --- | --- | --- |
|  | Mean | SD | Mean | SD | Mean | SD |
| 5 | 712.49 | 153.23 | 3316.17 | 378.14 | 4.76 | 0.89 |
| 25 | **229.06** | 64.05 | **876.42** | 168.02 | **4.00** | 1.11 |
| 60 | **39.39** | 4.44 | **131.68** | 22.69 | **3.35** | 0.45 |

The data was collected following a single intravenous administration to male Sprague Dawley rats (1 mg/kg); LLOQ: 5.10 ng/mL for plasma and 2.04 ng/mL for brain.

**Figure 5.**
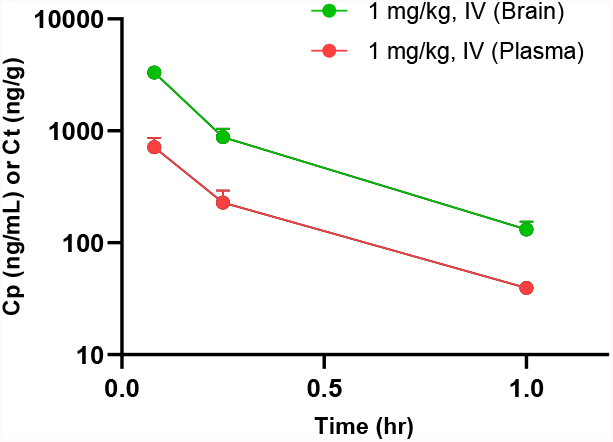
Plasma and brain concentration-time profiles (mean ± SD) of compound **26**.

### Radiochemistry and lipophilicity

[^11^C]**26** was synthesized in excellent radiochemical purity (> 99%) and high molar activities (170 - 220 GBq/µmol) with radiochemical yields of 34 ± 7% (n = 12) relative to [^11^C]CO_2_ at the end of bombardment (Figure 6A). [^11^C]**26** was reformulated in saline with less than 10% ethanol (v/v) as an injection solution. At three time points (30, 60, 90 min), the *in vitro* stability was tested, and no obvious degradation or radiolysis was found, which is shown in Figure 6B-C. The lipophilicity of [^11^C]**26** was evaluated by the ‘shake-flask’ method, revealing a Log*D*_7.4_ valve of 3.67 ± 0.04 (n = 3).

**Figure 6.**
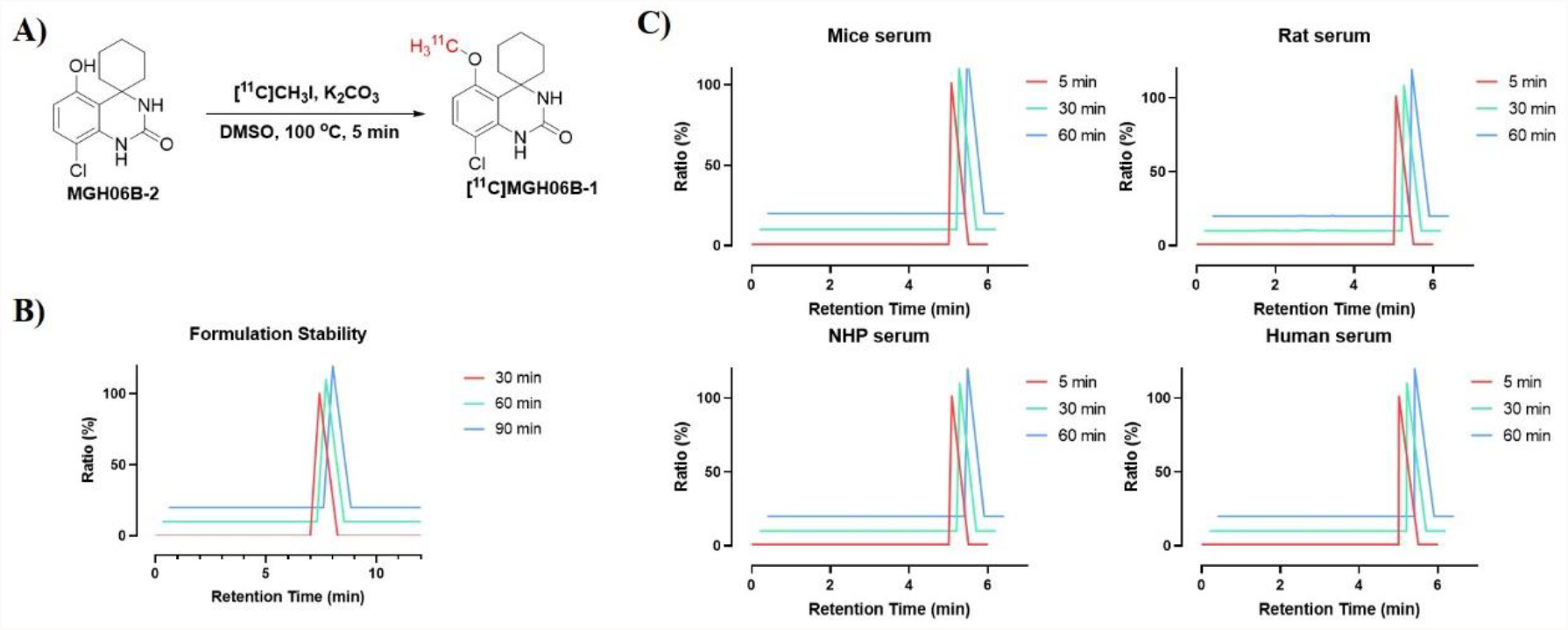
Carbon-11 labeling of [^11^C]**26** and in vitro stability studies. (A) Carbon-11 labeling conditions; (B) stability in PBS (containing 10% Ethanol, v/v); (C) stability in serum of various species (rodents, NHP and human).

### Plasma protein binding and *in vitro* cell uptake

The plasma protein binding was determined in mouse, rat, and human plasma, respectively. The results are presented as free fraction ratio in Figure 7A, which demonstrated a reasonable free fraction (≥1%) of **26** in the plasma. *In vitro* cell uptake assay indicated that the radioactivity uptake of [^11^C]**26** was 6-7 folds higher in HEK293-PDE7B recombinant cell lines at 30- and 60-min incubation time, respectively (Figure 7B). The blocking study of [^11^C]**26** was carried out at 60 min incubation time with unlabeled compound **26** (1 μM) and demonstrated high specific binding (ca. 90% blockade) in cell uptake experiments (Figure 7C).

**Figure 7.**
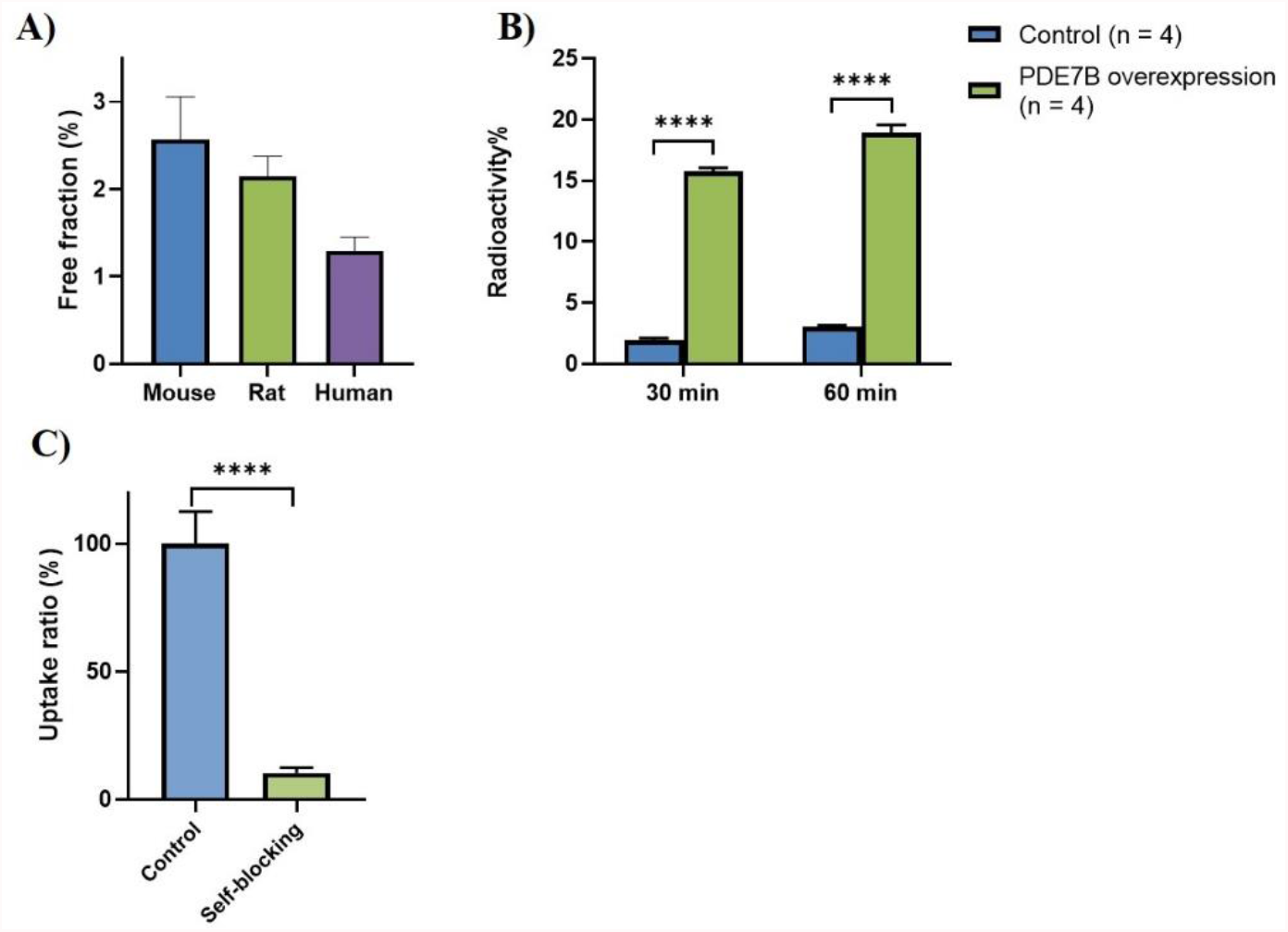
A) Protein binding potency in mouse, rat, and human plasma; B) Radioactivity uptake comparison between control and PDE7B over-expressed cells; C) Blocking experiments in PDE7B over-expressed HEK293 cells using compound **26**.

### PET imaging in rodents

[^11^C]**26** was injected into mice for microPET studies (1.85 MBq per mouse and 52-53 MBq per rat; molar activity 170 - 220 GBq/µmol). Time-activity curves (TACs) demonstrated that the tracer rapidly accumulated in the brain, whereas peak SUV was reached within 2 min p.i. (see supporting information, Figure S1). Similarly, baseline and self-blocking PET experiments at a dose of 1mg/kg were conducted in SD rats to confirm *in vivo* specificity of [^11^C]**26**. In rats, the highest SUV values (>2) were reached at approximately 5 min post injection, followed by a washout phase, ultimately reaching a plateau of 0.5 (SUV) at 20 min post injection (Figure 8). [^11^C]**26** showed heterogeneous brain uptake, with the highest tracer uptake observed in the striatum and cortex, followed by the cerebellum in the initial 10 min post injection. Notably, however, tracer washout from the cerebellum was substantially slower than in other brain regions. Pretreatment with non-radioactive compound **26** only showed marginally reduced brain uptake (see supporting information, Figure S2).

**Figure 8.**
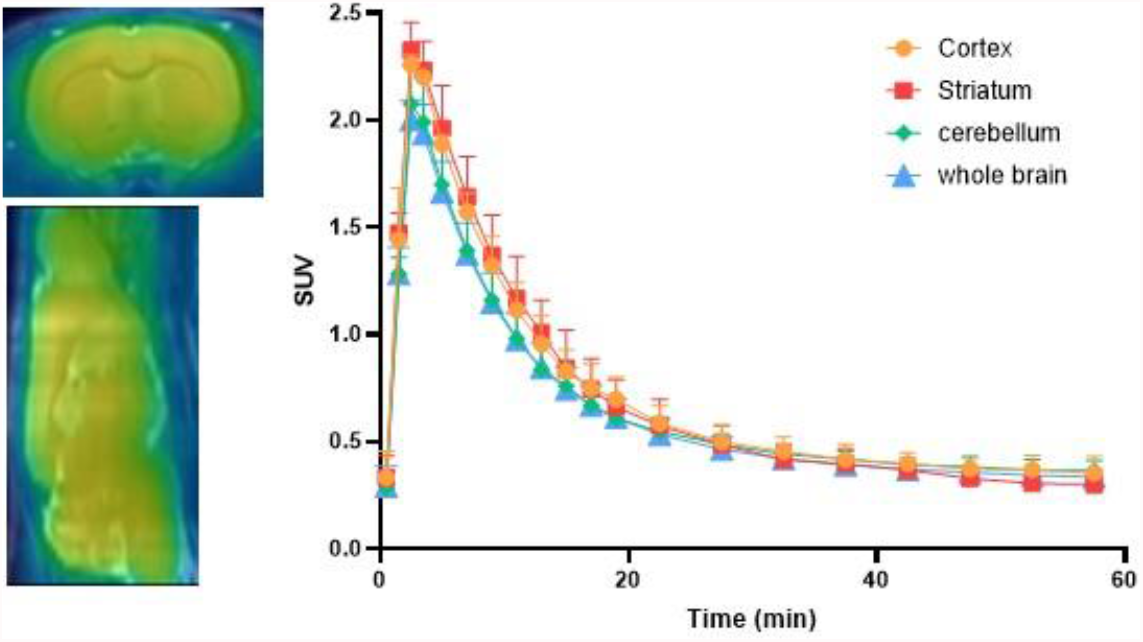
Summed PET images (0-20 min) in rat brain following injection of [^11^C]compound **26** and the corresponding TACs of the cortex, striatum, cerebellum, and whole brain from 0 to 60 min.

### PET imaging in nonhuman primates

[^11^C]**26** (200-278 MBq) was injected into monkeys under baseline and self-blocking conditions, and summed PET images are presented in Figure 9. TACs indicated that [^11^C]**26** rapidly crossed the BBB and reached the peak SUV (SUV_max_ = 3) at ca. 3 min p.i. Consistent with rat data, high tracer uptake was observed in PDE7B-rich monkey brain regions, including the cortex, putamen, caudate, followed by relatively lower uptake in the cerebellum (Figure 9A). The radioactivity uptake in all assessed regions was moderately reduced by pretreatment with non-radioactive **26** at the dose of 0.4 mg/kg or 1 mg/kg, respectively (Figure 9B).

**Figure 9.**
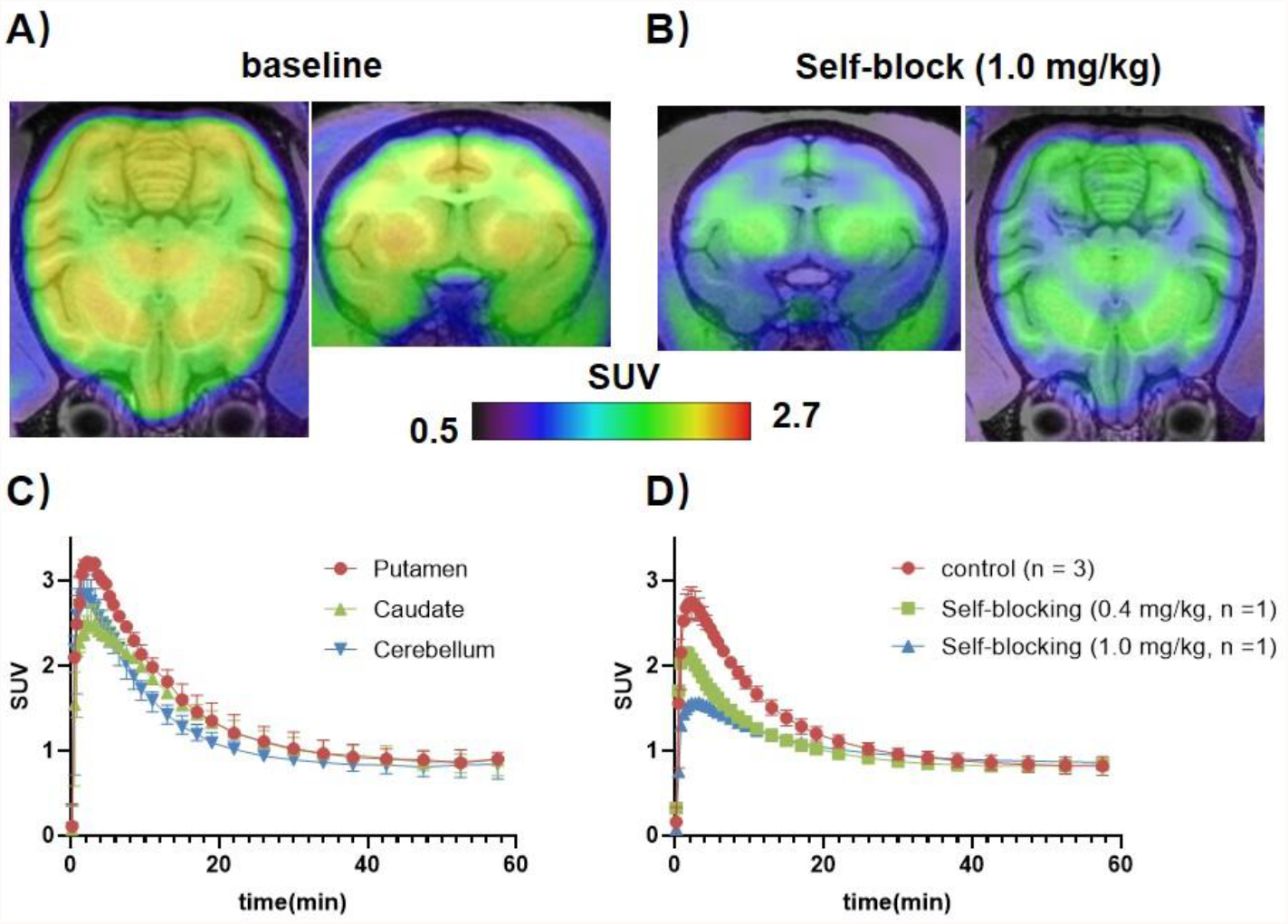
PET images and TACs in the monkey brain following injection of [^11^C]**26**. (A) Representative whole-body PET images of the baseline scans, averaged from 0-20 min; (B) Whole-body PET images under self-blocking conditions (1.0 mg/kg), averaged at 0-20 min; (C) Whole brain TACs of the baseline scan from 0 to 60 min and (D) Whole brain TACs at different blocking doses.

### Whole-body *ex vivo* biodistribution studies in mice

To study the distribution of [^11^C]**26** in the whole body, *ex vivo* biodistribution analysis was carried out in CD-1 mice at four different time points (5, 15, 30, and 60 min post injection, Figure 10). At 5 min p.i., the radioactivity was highly accumulated in the kidney, liver, brain, muscle, and particularly in the heart (up to 24% ID/g). Although initial cardiac uptake was the highest, it decreased to less than 2% within 60 min. The radioactivity was rapidly washed out from the blood, spleen, lung, pancreas, stomach, and small intestine. In contrast, high retention in the liver was found at 60 min post injection, suggesting a hepatobiliary clearance pathway.

**Figure 10.**
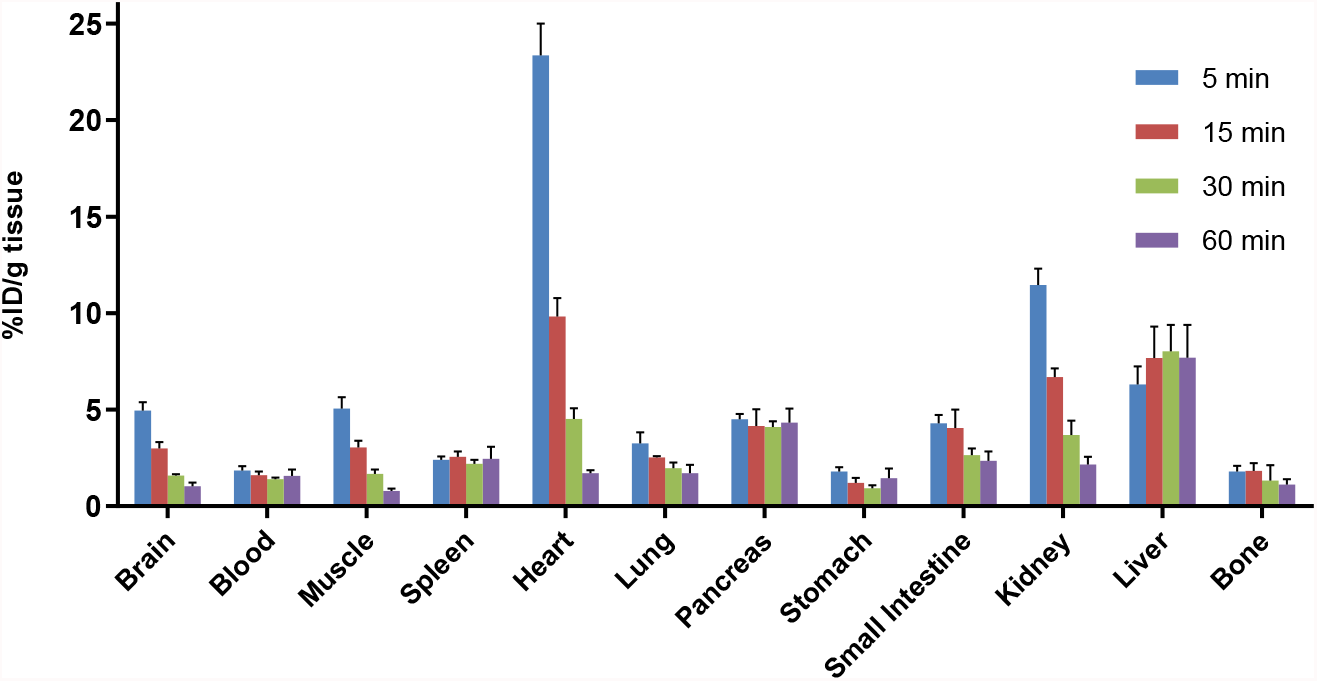
*Ex vivo* whole body biodistribution in CD-1 mice at four different time points (5, 15, 30 and 60 min) post injection of [^11^C]**26**. The results are expressed as the percentage of the injected dose per gram of wet tissue (% ID/g).

### Radiometabolite Analysis

Radiometabolite studies of [^11^C]**26** in rat brain and plasma (n = 2) were performed to evaluate *in vivo* stability (Figure 11). More than 98% intact [^11^C]**26** was found in the rat brain at 5 min. At 30 min post injection, assessment of brain homogenates still showed more than 75% of the parent fraction. Compared to that of the brain, the clearance of [^11^C]**26** in plasma was substantially faster with 70% ±18% and 19% ± 0.6% parent fraction found in the plasma at 5 min and 30 min p. i., respectively.

**Figure 11.**
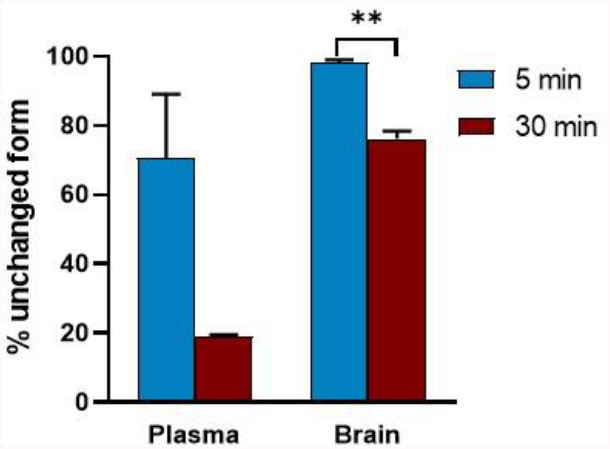
Radiometabolite study of [^11^C]26 in rat brains and plasma (n = 2) at the time point of 5 and 30 min post injection

## Discussion

In the present study, we report on the preclinical development of [^11^C]**26** as a potential PDE7-targeted PET radioligand. We were able to establish a simple and efficient synthetic route that allowed the synthesis of both reference compound and precursor in appropriate quality for radiochemistry and further biological assessment. Comparison between IC_50_ values of **26** towards PDE7 and other PDE isoenzymes demonstrated that compound **26** was highly potent and selective for PDE7. The former was supported by molecular docking analysis, unveiling significant interactions between **26** and several amino acids at the hydrophobic binding pocket of PDE7. ADME parameters, including plasma protein binding, CYP450 profiling, and hERG channel assay indicated a beneficial safety profile and NeuroPK studies in SD rats revealed that the brain-to-plasma ratio was >3 at all measured time points, indicating a high likelihood of brain penetration.

[^11^C]**26** was efficiently synthesized from its corresponding phenolic precursor **27** in one step using [^11^C]CH_3_I, providing high radiochemical yields and high molar activity. Further, PDE7B overexpressing HEK293 cells^8, 9^ were used to corroborate whether [^11^C]**26** binds to the PDE7B isoform in vitro. [^11^C]**26** showed substantially increased cell uptake in PDE7B overexpressing versus control cells, whereas the high *in vitro* specificity was confirmed by blocking studies.

*In vivo* dynamic PET scans of [^11^C]**26** across different species (mice, rats, and monkeys) confirmed the anticipated brain penetration of this ligand. After intravenous administration, the radioactivity quickly permeated through the blood-brain barrier and distributed heterogeneously in the brain. In rodents, tracer uptake was slightly higher in the striatum and cortex than that of the cerebellum, particularly in the initial 10 min post injection (Figure 8). Interestingly, there was a slower tracer washout from the cerebellum, as compared to all other brain regions, indicating that there might be a distinct mechanism of retention in this particular region in rats. While initial tracer uptake was significantly higher in the NHP cerebellum, as compared to the rat cerebellum, a faster tracer washout was observed from the NHP cerebellum. These results suggest that species differences in tracer kinetics are particularly pronounced in the cerebellum, and that caution is warranted with regard to the extrapolation of rodent finding to higher species in this particular region. In addition, a dose-dependent signal reduction upon blocking experiments with non-radioactive **26** in the nonhuman primate (NHP) brain suggested that the tracer may be used to assess target occupancy in higher species (Figure 9). The results of whole-body *ex vivo* biodistribution in CD-1 mice were consistent with PET findings. In consideration of the rapid washout from the kidneys and the sustained retention in the liver over 60 min (Figure 10), the radioactivity may be excreted mainly via the hepatobiliary pathway. Indeed, although radiometabolite studies of [^11^C]**26** demonstrated high *in vivo* stability in the brain, radiometabolites were detected at 30 min post injection in the plasma, which may explain the high liver uptake at later time points. Overall, we conclude that [^11^C]**26** shows promising performance characteristics in vitro; however, only limited in vivo specificity. Further structural modifications, leading to an improved inhibitory constant in the single-digit nanomolar range, may improve the degree of specific binding *in vivo*.

## Conclusion

This work describes the *in vitro* and *in vivo* evaluation of PDE7-targeted PET radioligand, [^11^C]**26** (codename [^11^C]P7-2104), which is endowed with a promising chemical scaffold of spiro cyclohexane-1,4’-quinazolinone. While target compound **26** proved to be potent and selective *in vitro*, cross-species *in vivo* studies with the radiolabeled analog, [^11^C]**26**, showed only moderate specific binding, thus warranting further structural modifications to achieve high *in vivo* specificity. The availability of suitable PDE7 PET radioligands is of critical value to understand the role of PDE7 in various disease pathologies as well as to guide preclinical and clinical drug discovery efforts.

## Acknowledgment

We thank the Division of Nuclear Medicine and Molecular Imaging, Radiology, MGH and Harvard Medical School, USA for general support. AH was supported by the Swiss National Science Foundation (SNSF).

## Notes

### Competing Interest Statement

The authors have declared no competing interest.

